# Ictal recruitment of anterior nucleus of thalamus in human focal epilepsy

**DOI:** 10.1101/788422

**Authors:** Emilia Toth, Ganne Chaitanya, Diana Pizarro, Sachin S Kumar, Adeel Ilyas, Andrew Romeo, Kristen Riley, Ioannis Vlachos, Olivier David, Karthi Balasubramanian, Sandipan Pati

**Affiliations:** Department of Neurology, University of Alabama at Birmingham, AL; Epilepsy and Cognitive Neurophysiology Laboratory, University of Alabama at Birmingham, AL; Department of Neurosurgery, University of Alabama at Birmingham, AL; Centre for Computational Engineering and Networking (CEN), Amrita School of Engineering, Coimbatore, Amrita Vishwa Vidyapeetham, India; Department of Electrical and Computer Engineering, Aristotle University of Thessaloniki, Greece; Brain Stimulation and Systems Neuroscience, Grenoble Institute of Neuroscience; Department of Electronics and Communication Engineering, Amrita School of Engineering, Coimbatore, Amrita Vishwa Vidyapeetham, India

**Keywords:** Thalamus, Discrete Wavelet Transform, Relative Wavelet Energy, Multiscale Spectral Entropy, Line Length, Epileptogenecity Index

## Abstract

The thalamic nuclei play diverse roles in the initiation, propagation, and termination of temporal lobe seizures. The role of the anterior nucleus of the thalamus (ANT) - a node that is integral to the limbic network is unclear. The objective of this study was to characterize temporal and - spectral patterns of ANT ictal recruitment in drug-resistant temporal lobe epilepsy (TLE). We hypothesized that seizures localized to the temporolimbic network are likely to recruit ANT, and the odds of recruitment were higher in seizures that had altered consciousness. Ten patients undergoing stereo-electroencephalography (SEEG) were recruited prospectively to record field potentials from the ANT. Using epileptogenicity index and line length, we computed the number of seizures that recruited the ANT (recruitment ratio), the recruitment latencies between the ANT and the epileptogenic zone (EZ), and latency of ANT recruitment to clinical manifestation for seventy-nine seizures. We observed that seizures localized to mesial temporal subregions (hippocampus, amygdala, anterior cingulate) have a higher predilection for ANT recruitment, and the recruitment was faster (ranged 5-12 secs) and preceded clinical onset for seizures that impaired consciousness. Seizures that recruited ANT lasted significantly longer (t=1.795, p=0.005). Recruitment latency was inversely correlated to seizure duration (r=-0.78, p=0.004). Electrical stimulation of the EZ induced seizures, in which early recruitment of ANT was confirmed. Stimulation of ANT did not induce a seizure. Finally, we tested the hypothesis that spectral and entropy-based features extracted from thalamic field potentials can distinguish its state of ictal recruitment from other interictal states (including awake, sleep). For this, we employed classification machine learning that discriminated thalamic ictal state from other interictal states with high accuracy (92.8%) and precision (93.1%). Among the features, the emergence of the theta rhythm (4-8 Hz) maximally discriminated the endogenous ictal state from other interictal states of vigilance. These results prompt a mechanistic role for the ANT in the early organization and sustaining of seizures, and the possibility to serve as a target for therapeutic closed-loop stimulation in TLE.

## Introduction

Temporal lobe epilepsy (TLE) syndrome is a highly prevalent focal epilepsy that can be surgically remediable if medications fail to control seizures(McIntosh *et al*., 2004). However, despite the decade of technological advancements, the surgical outcomes remain suboptimal(Thom *et al*., 2010). While delineating the seizure focus continues to be the central dogma of optimizing surgical and neuromodulation therapies, growing evidence has implicated TLE as a network disorder with epileptogenic “focus” extending beyond the hippocampus to a network of cortical and subcortical structures including the thalamus (Spencer, 2002; Keller *et al*., 2015; He *et al*., 2017). Imaging studies have identified thalamic hubness as a predictor of poor surgical outcome and persistence of thalamo-temporal connectivity post temporal lobectomy is correlated with seizure recurrence (Sperling *et al*., 1990; Newberg *et al*., 2000; He *et al*., 2015; He *et al*., 2017). The corticothalamic projection provides massive top-down input to the thalamus, providing the anatomical connectivity for seizure propagation, while the reciprocal thalamocortical projections regulate information flow to the cortex, thereby providing a physiological basis for modulating ictogenesis (Guye *et al*., 2006; Huguenard and McCormick, 2007). The thalamus is not a unitary structure and comprises of multiple nuclei that exhibit variable cortical connectivity(Hwang *et al*., 2017). Experimental studies suggest that the thalamic nuclei play diverse roles in initiation, propagation, and termination (collectively termed as ictogenesis) of limbic seizures(Bertram *et al*., 2001; Takebayashi *et al*., 2007; Langlois *et al*., 2010; Feng *et al*., 2017). In TLE, prior clinical studies have explored the influence of lateral and medial thalamic nuclei in ictogenesis, but none to date have studied the anterior nucleus of the thalamus (ANT) - a node that is integral to the limbic network(Child and Benarroch, 2013; Evangelista *et al*., 2015; Romeo *et al*., 2019).

The ANT is proposed to propagate and sustain focal limbic seizures and has been targeted for neuromodulation therapy in epilepsy(Hamani *et al*., 2004; Fisher *et al*., 2010; Salanova *et al*., 2015) Chemical inactivation (muscimol to inhibit neuronal firing)(Bittencourt *et al*., 2010), lesioning(Takebayashi *et al*., 2007) or high-frequency electrical stimulation of ANT in preclinical models disrupted the progression of convulsive seizures to bilateral tonic-clonic seizures(Takebayashi *et al*., 2007; Hamani *et al*., 2008). Preliminary clinical studies from our group and others have demonstrated that neural activity within the ANT is increased significantly at the onset and early propagation of temporal lobe seizures(Osorio *et al*., 2015; Pizarro *et al*., In-Press). In the present study, we have characterized the temporal trends and patterns of ictal recruitment of ANT in consenting adults undergoing stereoelectroencephalography (SEEG) for localization of suspected mesial TLE. We hypothesized that seizures localized to the temporolimbic network are likely to recruit ANT, and the odds of recruitment were higher in seizures that altered consciousness. Thus, at first, we identified the epileptogenic zone (EZ) for all the studied seizures using a validated quantitative metric called the “Epileptogenicity Index (EI)”(Bartolomei *et al*., 2008; Roehri *et al*., 2018). From this, we selected the seizures that had temporolimbic subregions as the EZ. The epileptogenic zone (EZ), as viewed by Bancaud and Talairach, is the site of the initiation and early propagation of the seizure (Talairach and Bancaud, 1966). Second, using an established seizure detection algorithm (line length)(Esteller *et al*., 2004) that was complemented with visual inspection, we confirmed the ictal recruitment of ANT and compared the detection latencies between the –a) EZ and ANT, and b) the ANT and clinical onset (behavioral manifestation) of temporal lobe seizures. Third, using demographic and other clinical covariables, we have estimated the predictors of ANT ictal recruitment in TLE. Finally, we tested the hypothesis that spectral and entropy-based features extracted from thalamic field potentials can distinguish its state of ictal recruitment from other interictal states (including awake, sleep). For this, we have used machine learning to evaluate the performance of spectral and temporal features in classifying ictal from interictal states. Prior studies have demonstrated that states of vigilance (SOV) can be distinguished purely using entropy or time-frequency (spectral) decomposition of EEG signals (Miskovic *et al*., 2019). In sum, using qualitative and quantitative metrics, we have characterized the recruitment of ANT in drug-resistant TLE syndrome.

## Materials and Methods

### Seelection of patients

All patients with suspected drug-resistant TLE undergoing SEEG exploration for mapping the EZ were eligible to participate in this study. The rationale for selecting TLE was based on its -a) structural connectivity to the ANT (Child and Benarroch, 2013); b) preclinical studies demonstrating recruitment of ANT in TLE(Hamani *et al*., 2004; Hamani *et al*., 2008; Sherdil *et al*., 2019); and c) the potentials of translating the knowledge gained in the study towards the development of a closed-loop DBS therapy. Before the surgery, written informed consent was obtained to implant and record from the thalamus for research purpose. The SEEG was clinically indicated in drug-resistant epilepsy with negative MRI (nonlesional) or where localization was inadequate for resective surgery from noninvasive investigations. The trajectory of one of the depth electrodes that was planned to target insula and operculum regions (for clinical purposes) was advanced further to sample from the ANT(Figure 2A). The strategy mitigated the increased risk and cost associated with the implantation of an additional depth electrode used exclusively for research. Clinical characteristics of the patients (N=10), including details of the pre-implant investigations, are detailed in (Table S1). The Institutional Review Board of the University of Alabama at Birmingham approved the study (IRB-170323005).

**Figure 1.**
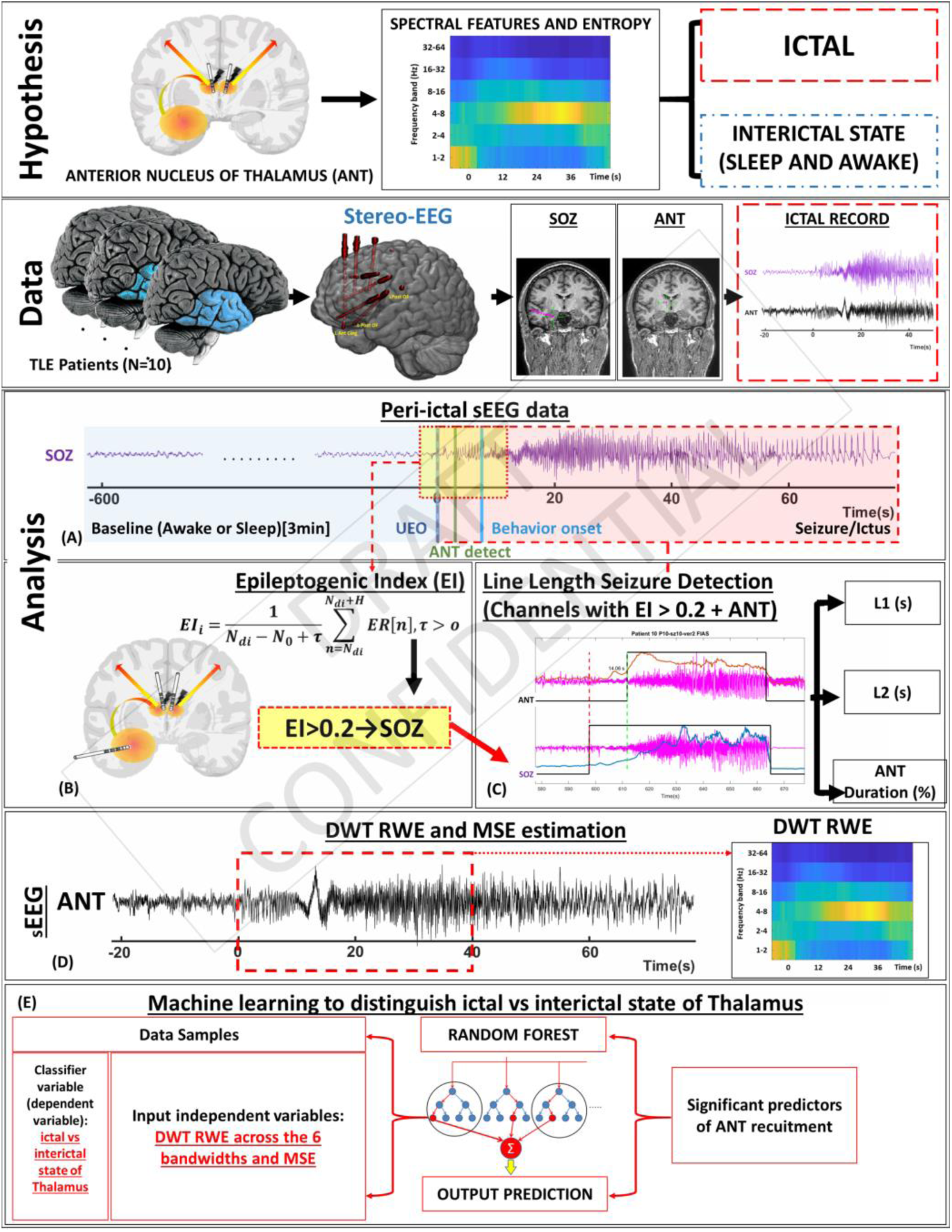
Methods Pipeline. The study aimed at testing whether the spectral and entropy features of the thalamic stereo EEG(sEEG) recording can reliably classify ictal state from interictal baseline activity observed during background sleep and wakefulness. Ten patients were recruited and underwent robotic sEEG implantation where the electrodes were consistently placed in 2 regions, the anterior nucleus of thalamus (ANT) and the clinician defined seizure onset zone (SOZ), along with other specific brain regions tailored to patient’s epilepsy. (A) Of the continuous monitoring, the sEEG was segmented from -600s before the electrographic onset (as detected by line length-LL) till the termination of the seizure. (B) Epileptogenicity Index (EI) was calculated on the sEEG data based on energy spectral density ratio (ER) ranging from 4-12Hz in lower frequencies and 12-127Hz in the higher frequencies. (C) LL seizure detection was performed on all channels which had an EI>0.2, including the ANT channels. (D) Based on the onset detected by the LL, the 6-level discrete wavelet transform relative wavelet energy (DWT-RWE) and Multiscale entropy (MSE) was calculated for the first 20sec after seizure onset and 180 seconds baseline data were extracted as features for machine learning. (E) Random forest machine learning was performed to test if the DWT-RWE (of the 6 frequency band widths) and MSE features estimated in the thalamus could reliably predict the ictal state vs the interictal baseline activity (during wakefulness and sleep).

**Figure 2:**
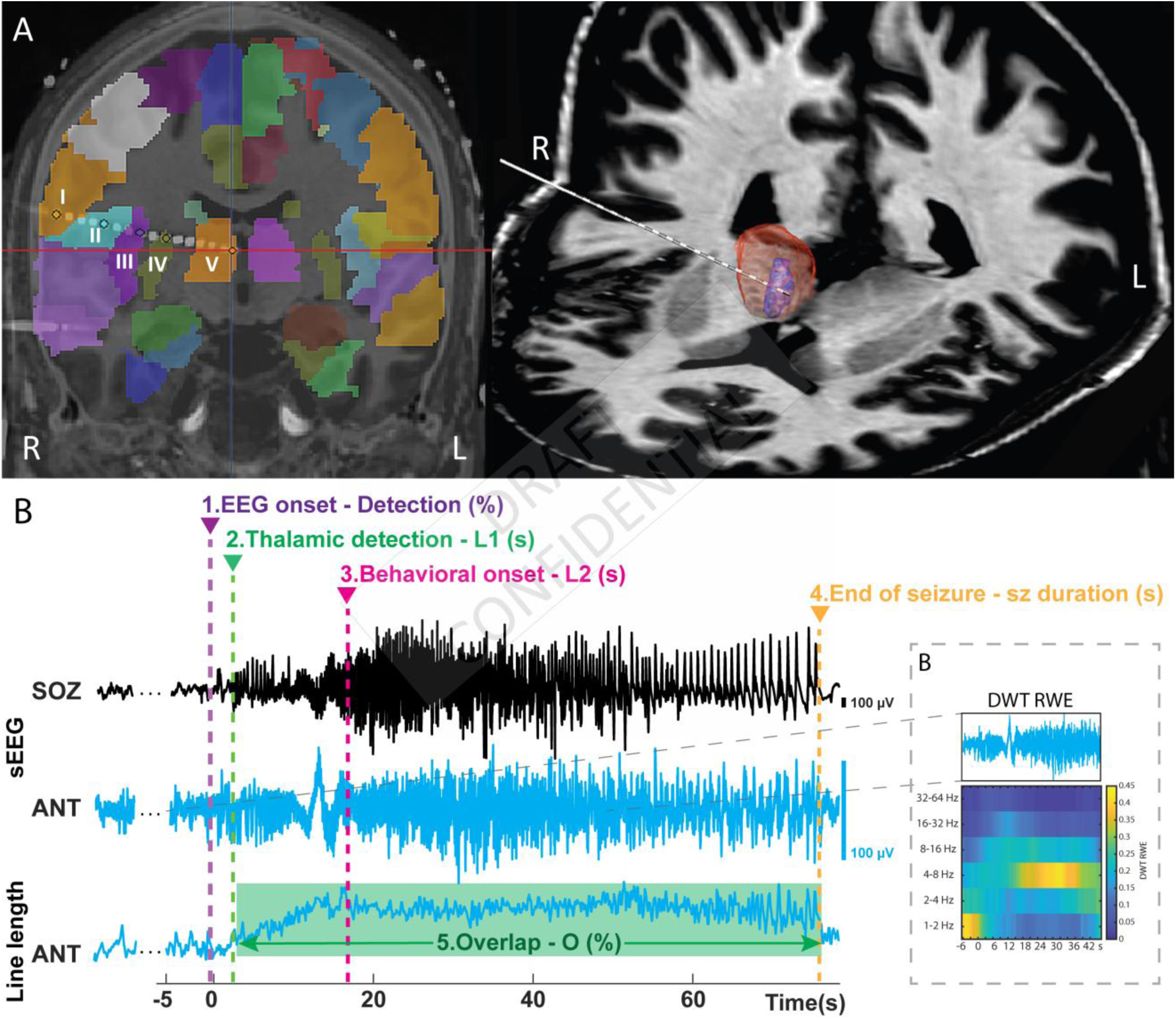
Electrode localization and Line Length (LL) detection, timing of seizure activity and time frequency analysis: (A) The analysis starts with identifying the unequivocal electrographic onset (UEO) by the clinician in the seizure onset channels (SOZ). LL was performed on all the cortical channels with EI>0.2 that represented the EZ and the 3 bipolar derivatives (i.e., linearly derived from 4 contact points) in the ANT. LL helps detect the ictal activity objectively. A seizure in ANT was detected when LL detected activity within 120sec of onset in SOZ, and if the activity lasted longer than 10s. Among the seizures detected in ANT, two latencies were estimated using LL. The first latency (L1) is the difference in LL detected seizure in the EZ and ANT while the second latency (L2) is the difference between clinical onset (behavioral manifestation) and LL detected seizure in the ANT. The duration of ANT recruitment as a percentage of the seizure duration in the EZ was termed as overlapped detection (O). (B) The discrete wavelet transform relative wavelet energy (DWT RWE) and multiscale entropy (MSE) were estimated on the LL detected ANT data.

### SEEG recording

Robotic assistance (ROSA device, Medtech, Syracuse, NY) was used to plan optimal and safe trajectories for SEEG multi-contact electrode implantation (12-16 contacts per depth electrode, 2mm contact length, 0.8mm contact diameter, 1.5mm inter-contact distance, PMT^®^ Corporation, Chanhassen, MN) to simultaneously sample both the ANT and pre-ordained temporal network regions of interest. Intracranial video-EEG was recorded using Natus Quantum (Natus Medical Incorporated, Pleasanton, CA, sampling rate 2048Hz) (Figure 1). Signals were referenced to a common extracranial electrode placed posteriorly in the occiput near the hairline.

### Reconstruction of depth electrodes into the brain

The following imaging sequences were used to confirm localization of electrodes: pre-implantation MRI sagittal T1-weighted images acquired on a 3T Philips Achieva (voxel size 1×1×1mm with a 170×256×256mm FOV) and post-implantation CT axial images acquired on a Philips Brilliance 64 scan (1mm slices with an in-plane resolution of 0.44×0.44mm with a 228×228×265mm FOV). It was prudent to establish that the implanted electrodes were in the ANT. Electrodes were localized using Lead-DBS software (Horn and Kuhn, 2015; Horn *et al*., 2019) (www.lead-dbs.org) and the trajectories were mapped using iElectrodes software (Blenkmann *et al*., 2017). Briefly, preoperative and postoperative patients’ images were linearly co-registered using Advanced Normalization Tools (Avants *et al*., 2011) followed by refinement with brain shift correction to improve the registration of subcortical structures (Schonecker *et al*., 2009). Both the images were normalized to ICBM 2009b NLIN asymmetric space using the symmetric diffeomorphic image registration. The data were then visualized using the Morel’s Thalamic atlas (Krauth *et al*., 2010). To track the trajectory of the electrodes, the normalized images were imported into iElectrodes, and the Automated Anatomic Labelling atlas (AAL) was overlaid to identify the structures through which the electrodes traversed (Figure 2A).

### Visual analysis of seizures in cortico-thalamogram

Seizure onset and offset were defined using standard criteria. Seizure onset was defined as the earliest occurrence of rhythmic or repetitive spikes in the cortex that was distinctive from the background activity, and that evolved in frequency and morphology. The EEG onset of the seizure in the cortex was marked as “unequivocal EEG onset” (UEO). Intracranial EEG segments were clipped such that these included 10 minutes (min) before the UEO marking and 10min post-termination of the seizure. The clipped video-EEG was annotated for: i) unequivocal behavioral changes that were defined as clinical onset, and ii) SOV (sleep or awake) before the onset of the seizure (UEO). SOV was classified as wakefulness or sleep (staging not performed) based on video and EEG (intracranial and simultaneously recorded limited scalp electrodes). In participants who have reported aura, the clinical onset time was determined as the time patient notified or pressed the event button. Seizures were classified based on semiology into focal aware seizures =FAS, focal impaired awareness seizures = FIAS and FBTCS= focal to bilateral tonic-clonic seizures (Fisher *et al*., 2017). A seizure without any clinical correlate but ictal electrographic activity lasting >10 seconds (s) was classified as electrographic seizures (ES)(Jirsch and Hirsch, 2007). Patients were assessed during and after the seizure by bedside nurses and EEG technicians for motor function, orientation (to time, place and person), speech (through conversation, naming objects and reading) and memory (by asking to remember an object and recollect after the seizure). These are standard protocols practiced in our epilepsy monitoring unit. A minimum of three seizures selected randomly among each seizure types (ES, FAS, FIAS, FBTCS) per patient, and overall 7-10 seizures per patient were included for analyses in the study. If a particular seizure type was less than three, we have included a higher number of the prevalent seizures to increase the overall N. Seizures with significant EEG artifact, or unavailable videos were excluded from the study.

### Quantification of the EZ

The epileptogenicity index (EI)(Bartolomei *et al*., 2008) has been shown to quantify the epileptogenicity of brain structures and has been validated with clinically identified probable EZ, seizure outcome following surgical resection, and with interictal high-frequency oscillation maps to identify EZ (Roehri *et al*., 2018; Vaugier *et al*., 2018). The EI statistically summarizes the spectral and temporal parameters of SEEG signals and relates to the propensity of a brain area to generate low voltage fast discharges. To compute EI, we have used the open-source software AnyWave (Colombet *et al*., 2015). Briefly, the energy spectral density ratio (ER) was estimated as a measure of abrupt increase in fast oscillations in the sEEG signal as a marker of seizure related change:

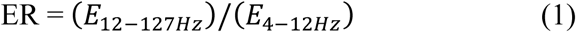

Use of Page and Hinkley’s cumulative sum algorithm helped optimize the time point of detection of fast oscillaitons. Finally, EI was calculated as the averaged ER over time just after detection of the rapid discharge in the first channel divided by the delay of involvement across other channels (Bartolomei *et al*., 2008). EI values from all the available seizures were computed from selected bipolar channels (first 3 channels from every electrode and the clinically noted channels) over the first 12sec of the seizure (starting 2sec before, and 10 s after UEO) and the highest EI value for any given channel across the seizures was used. Channels demonstrating EI value above 0.2 were considered within the EZ (Roehri *et al*., 2018). We ensured that the EZ overlapped with the clinically identified SOZ.

### Seizure detection algorithm

Line length (LL) was used to detect the ictal activity objectively. The LL feature was derived as a simplification of the running fractal dimension of a signal (Esteller *et al*., 2004). It measures the length of the signal in a particular window and compares it to a threshold. The length of the signal is dependent on the amplitude and frequency of the signal, making this feature highly suitable to sense changes in amplitude and/or frequency that typically occur during seizures. The change in LL was considered significant when the measured LL in that segment was greater than 2SD compared to the mean of a 3min baseline segment for at least 10sec. Preprocessing steps involved detrending within 2s, removing DC (Matlab detrend), reconfiguring in bipolar montage and removing 60 Hz line noise with a notch filter (2^nd^ order Butterworth zero phase shift filter – Matlab designfilt and filtfilt). Subsequently, the LL was calculated on this preprocessed data over 0.25s windows with 50% overlap as shifting windows and the resultant LL data was smoothed using the Matlab function ‘movmean’ over 2s windows. The LL was calculated for each segment with *m* samples using the following formula:

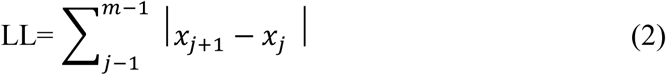

Where *m* is the number of sample points, and the *x*_*j*_ is the sum of absolute distances between consecutive *m* sample points.

The LL detections were merged if they were closer than 2s. When the merged detections were longer than 10s, they were considered by the detection algorithm as seizures (Pizarro *et al*., In-Press). To improve the accuracy of the seizure detection of the entire event, on the basis of the length of the seizure, another *layer* of temporal summation was performed wherein, if the10s-merged-detections were closer than 5s, they were again merged to constitute one single seizure event. The method helped to avoid detecting them erroneously as multiple close-lying seizures, thus helping in the detection of slower ictal spiking activity.

The electrophysiological features of ANT involvement during a seizure are not well described. Thus, we have adopted a manual offline seizure detection approach based on LL features described above. LL was performed on all the cortical channels with EI>0.2 that represented the EZ and the bipolar derivatives obtained from recording the ANT. The visually determined UEO in the cortex was used as the reference time stamp to mark seizure onset for that event. We then estimated another measure based on LL detection of seizure onset in each of the channels (i.e., based on the electrographic changes in the channels, LL could show a variation in the detection latency from one channel to another). Seizures in ANT were reported as “not detected (ND)” when the specified changes in LL as described above were – a) absent in the first 120s after LL detected seizure in the cortex (LL UEO); or b) present transiently (<10% of total seizure duration) in the ANT for the entire duration of the seizure.

### Defining parameters to estimate temporal trend-Recruitment ratio and latencies

We defined the term *recruitment ratio* as the % of seizures that were detected by LL in the ANT divided by the total number of seizures in that group(Figure 2B). Among the seizures detected in ANT, two latencies were estimated using LL. The detection latencies are estimated in reference to the LL detected seizure in the EZ (t=0 secs) (Figure 2B). The first latency (*L1/Recruitment Latency*) is the temporal difference in LL detected seizure between the ANT and EZ (positive values indicate detection in ANT succeeded detection in the EZ). The second latency (*L2/Behavioral latency*) is the difference between LL detected seizure in the ANT and clinical onset (behavioral manifestation) (Figure 2B). Negative L2 implies thalamic recruitment preceded the clinical onset. We also expressed the duration of ANT recruitment (termed as *overlap-O*) as a percent of the seizure duration in the EZ using the following formula:

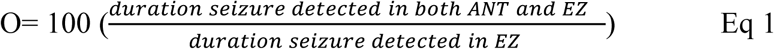

### Time-frequency decomposition of thalamogram

The time-frequency decomposition of field potentials was performed with the discrete wavelet transform relative wavelet energy (DWT –’db4’, RWE, 6 levels). The DWT provides a non-redundant, highly efficient wavelet representation and direct estimation of local energies at the different relevant scales (Ursino *et al*., 2004; Chen *et al*., 2017). The motivation for selecting DWT is based on prior studies suggesting this method can be an optimal tool for online seizure detection that can be translated in implantable neural prosthesis (Kamboh *et al*., 2007; Narasimhan *et al*., 2011; Logesparan *et al*., 2012; Yang *et al*., 2014). The DWT RWE was calculated on 4s windows, with 3 s overlap, for 1-2 Hz, 2-4 Hz, 4-8 Hz, 8-16 Hz, 16-32 Hz and 32-64 Hz from awake, sleep and seizure segments of the thalamic signal. Awake and sleep segments were at least 6min before the LL detected UEO while the seizure segments were 20s after the LL detected seizure in ANT.

### Multiscale entropy analysis of thalamogram

We calculated multiscale entropy (MSE) to characterize the temporal predictability of a time series across several time scales in the thalamus, serving as an index of its capacity for processing information (Miskovic *et al*., 2019). While estimating the MSE, the maximum template length (*M*) was set to 3, and the matching threshold was set to 0.5, scaled to 10. The data was averaged across the 10 scales to indicate the mean MSE for that 4s time window. MSE was calculated across the length of the ANT SEEG signal on moving window epochs (awake, sleep and seizure segments) like DWT analysis.

### Direct Electrical Stimulation of the epileptogenic cortex and the ANT

Direct electrical stimulation (DES) with an attempt to induce a seizure and confirm localization of EZ is an acceptable procedure that is performed routinely in our epilepsy center. The stimulation is performed typically towards the end of the evaluation after the recording of multiple spontaneous seizures and after restarting of antiseizure medications. Since DES of the thalamus is not a standard procedure, IRB approval and informed consent were obtained before stimulation. Stimulation parameters were square wave biphasic pulses, delivered in a bipolar montage with width 200-500 µs, frequency 10, 20 or 50 Hz, pulse duration 3-5 secs, and current ranged between 3-8 mA. The goals of the stimulation were two folds-a) to confirm thalamic recruitment for seizures induced from the EZ, and b) if stimulation of ANT can induce a habitual seizure.

### Statistical analysis

To examine the influence of clinical factors such as (1) seizure type and SOV on EI, L1, L2 and O (Figure 3A, 4E, 4F, 4G), and (2) impairment of consciousness on EI (Figure 3B), we used ANOVA, with Bonferroni correction. Pearson’s correlations coefficients were used to test the relationship between (1) seizure duration and recruitment latency (L1), (2) overlap (O) and seizure duration, (3) recruitment latency (L1) and behavioral latency (L2). The clinical predictors of ANT recruitment (ie., age at epilepsy onset (years), epilepsy duration (years), reported seizure frequency per month, seizure duration in seconds) were tested using logistic regression, and were controlled for gender, subject, history of status epilepticus and MRI lesion. Since DWT-RWE showed non-normal distribution, we used Wilcoxon rank-sum (WRS) test (Hollander and Pena, 2004) to test how the ictal spectral patterns differed (i.e., increase or decrease in DWT-RWE) from baseline (awake and sleep) state. Five DWT RWE samples of moving windows of 4s length, with 3s overlap were input into the WRS test to form time-related spectral power distribution subset from the seizures with ANT recruitment separated by seizure types (ES=7, FAS= 8, FIAS=24, FBTCS=6) for every frequency band (n=35, 40, 120, 30). These subset distributions were compared to awake and sleep interictal distribution (n=17040) *with Wilcoxon test*, and FDR correction (Genovese *et al*., 1997; Genovese *et al*., 2002; Nichols and Hayasaka, 2003; Delorme and Makeig, 2004) was applied on the resulting p values.

**Figure 3.**
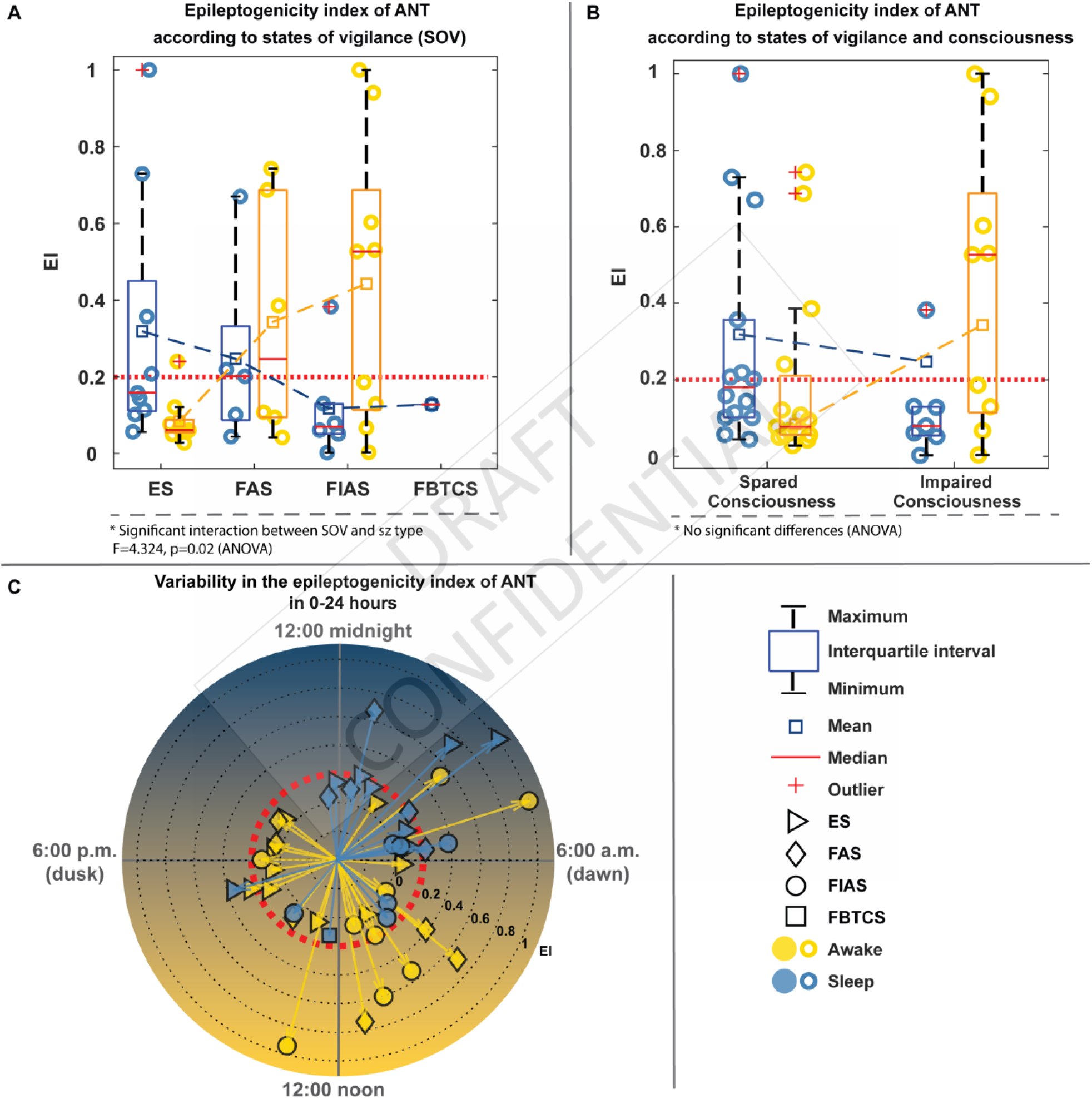
Epileptogenecity Index (EI) of Anterior Nucleus of Thalamus (ANT) and its clinical relationship: The three graphs represent the variability in EI estimated in the thalamus based on the clinical scenario in which it is tested. (A) ANOVA testing the effect of seizure type and SOV, shows no overall effect of either seizure type or SOV, but there is a cross over interaction showing the effect of seizure type on the EI varies depending on the SOV (f=4.32, p=0.02). (B) shows no significant difference in thalamic EI between those seizures that were associated with impaired consciousness and those without impaired consciousness and (C) is a clock-plot indicating the variability in thalamic EI over a 24hours period. As evident FIAS and FAS were clustered between 12 midnight to 12 noon and their respective thalamic EI values were higher too. The right lower panel is an index of the symbols used in the figure.

### Machine Learning with Random Forest and Random Kitchen Sink

Supervised machine learning (Random Forest) is used to classify the given samples (thalamic EEG) into two classes: seizure and interictal states (combined awake and sleep). Seven features namely: DWT RWE in 6 frequency bands and the average of the first 10 scales of the MSE of awake, sleep and 20 s seizure segments from the ANT were used as features for random forest and random kitchen sink machine learning algorithms. The seizure class contains 373 data points, and physiological class has 2850 data points (approximately 4 hours of data), each being represented by 7 features. The data is unbalanced, and hence the smaller data was split in 80%: 20% for training and testing data. 300 data points are chosen at random for training, and 73 data points are chosen for testing purposes from each class, and random forest (RF) classification technique is applied using all the features to calculate the accuracy, precision (positive predictive value) and recall (sensitivity) values. To ensure that training and testing are done in an unbiased manner, the process is repeated 500 times by choosing the training and test samples randomly during each iteration, and the average values of the accuracy, precision, and recall are used. The entire procedure is run using 5,10,15, and 20 estimators and four sets of results are presented that indicated that we have an improved classification with an increasing number of estimators(Breiman, 2001; Altmann *et al*., 2010).

In order to validate the results of one prediction model, we tested the same using Random kitchen sink (RKS) learning, which involves a non-linear mapping of the features on to a higher dimension space and enabling the features to become linearly separable in the higher dimension(Rahimi and Recht, 2009). The feature mapping is done using a real Gaussian function as the Radial basis function (RBF) Kernel. This, in combination with a regularized least square algorithm for regression, allow us to obtain a simple classifier that can be used for real-time applications.

### Data availability

The data that support this study are available from the corresponding author, upon reasonable request.

## RESULTS

### Seizure types and their localization

Seventy-nine seizures from 10 subjects were analyzed in this study (Clinico-demographic details in Table S1). None of the patients had any bleeding from the thalamic implant. Based on the consensus on anatomo-electroclinical features and EI, the EZ was determined to be mesial temporal (2), medial-lateral (2), temporal pole (1), temporal plus (2), and multifocal subtypes (3; including one bitemporal). Based on the EI values (>0.2), 66/79 seizures had EZ involving the hippocampus (H), amygdala (A) and/or anterior cingulate (AC) complex (termed HAAC complex). The remaining 13 seizures had EI based onsets involving temporal pole, anterior insula, orbitofrontal, posterior cingulate, and lateral superior temporal gyrus. We have classified this group as extra-HAAC seizures (Figure 4B). Among the HAAC seizures, 18 were ES, 15 were FAS, 27 were FIAS and 6 were FBTCS. Sixty-two percent (41/66 of the HAAC onset seizures were noted during sleep.

**Figure 4.**
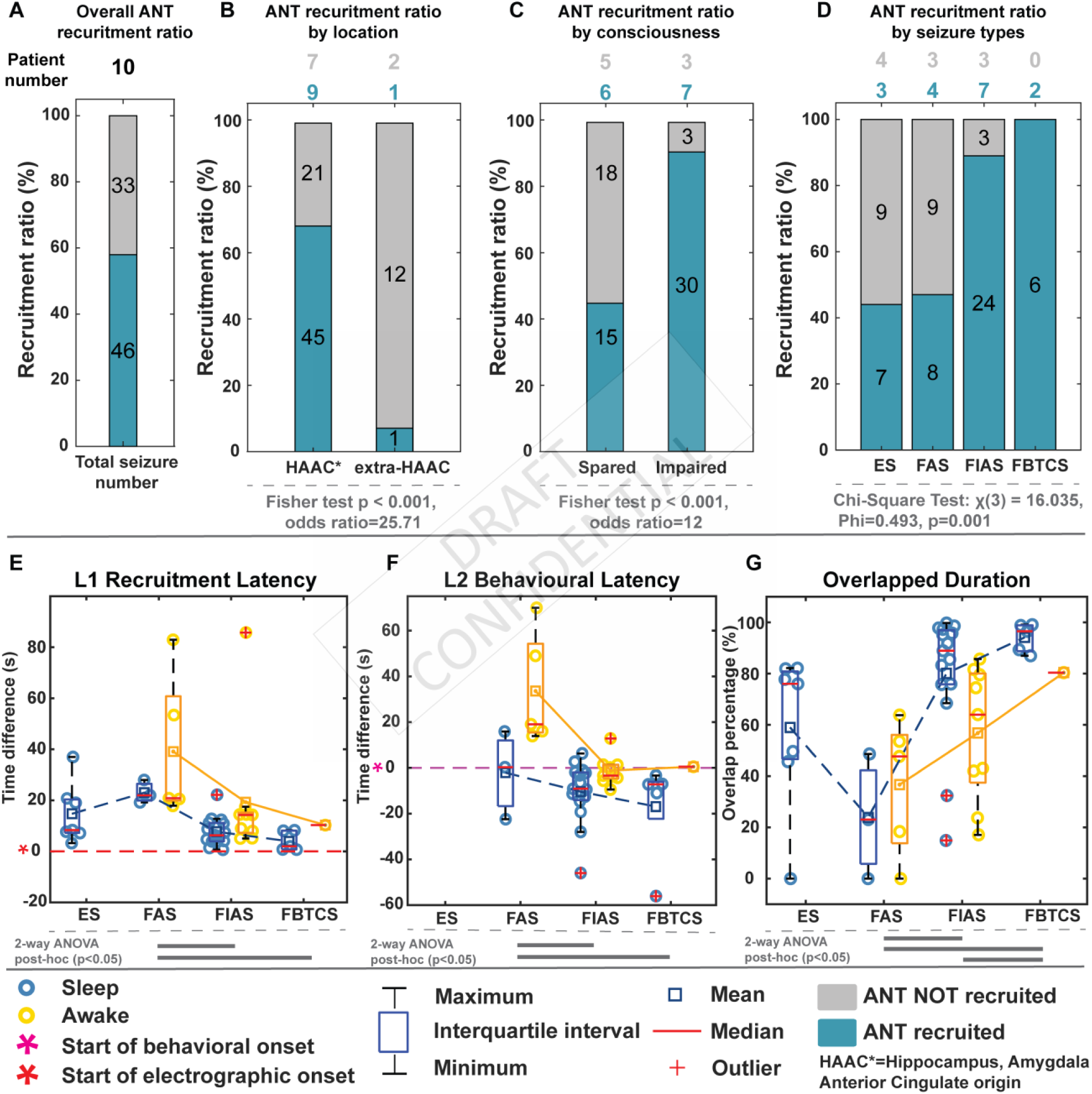
Clinical phenotypes of ictal recruitment of anterior nucleus of thalamus (ANT): (A) Overall 46 seizures of a total of 79 showed ictal thalamic recruitment, of which 45 seizures (B) had the seizure originating in the hippocampus, amygdala anterior cingulate complex (HAAC). (C) Among the seizures that originated in HAAC complex, we found that seizures with impaired consciousness had a higher incidence of thalamic ictal recruitment. The odds of thalamic recruitment was 12 times greater with impaired consciousness seizures (p<0.001). (E) L1 latency was found to be significantly different between the FAS vs FIAS (p=0.018) and FAS vs FBTCS (p=0.037). The main effect of SOV and the interaction between the SOV and seizure type were not significant. (F) However, L2 latency significantly different between the seizure types (f=5.6, p=0.008) and the SOV (f=8.23, p=0.007). The interaction between seizure type and the SOV was not significant. (G) ANT thalamic overlap with the duration of the seizure was different across the seizure types (f=18.36, p=0.002). The interaction between the seizure type and SOV was also significant (f=7.2, p=0.002).

### Epileptogenicity index of the anterior nucleus of the thalamus

The ANT EI was higher (>0.2) in 21% (n=16) of the seizures recorded from five subjects, suggesting that ANT was recruited early during seizure initiation and propagation (Table S2). Twelve seizures with higher ANT EI had their onset in the HAAC complex. The remaining seizures were from one subject #8 only, that had onset in posterior or mid-cingulate regions (Table S1 and S2). The higher ANT EI was not restricted to any particular seizure type or pre-seizure SOV (Figure 3 A-C). Though seizure type and SOV did not have distinctly different EIs, there was a cross over interaction i.e., EI of the different seizure types are dependent on the different SOV (f=4.32, p=0.02) (Figure 3A). Seizures with impaired consciousness did not show any difference in mean EI compared to consciousness-sparing seizures (Figure 3B).

### Temporal trends in ictal recruitment of anterior thalamus

The line length detected 59% of seizures (n=46 out of 79 seizures from 9 out of the 10 participants) in the ANT (Figure 4A). The only subject (#9) without any seizure detected by the LL in ANT had only two FAS towards the end of three weeks monitoring. One of the FAS had significant motion artifact that precluded analysis, but visual inspection showed ictal changes in the ANT. Out of the 46 seizures detected in the ANT, 45 of them had onset in the HAAC regions (Figure 4B). Only one seizure with an onset in the mid and posterior cingulate was detected in the ANT. ANT was recruited in 100% and 89% (n=24 of 27) for FBTCS and FIAS respectively (Figure 4D). The detection latencies in the ANT (L1) was found to be significantly different between the FAS (34s) vs FIAS (12.04s) (p=0.018) and FAS (34s) vs FBTCS (5s)(p=0.037)(Figure 4E). The ANT was recruited before the clinical/behavioral onset (L2), and the recruitment was faster during sleep than during wakefulness (Figure 4F). The mean latencies (L2) were: FAS +20.69 secs, FIAS -6.9 secs, FBTCS -14.01 secs (negative values indicate detection in the ANT preceded behavioral onset). Overall, the detection in the ANT preceded behavioral onset in 70% of FIAS and for all except one in FBTCS. The ictal electrographic changes persisted in the ANT for a variable duration (expressed as a percent of total seizure duration, overlap): 55% for electrographic seizures, 47% for FAS, 72% for FIAS, and 92% for FBTCS (Figure 4G).

### Correlations between recruitment latencies and seizure duration

Correlation analyses were performed between the temporal parameters of ANT recruitment and seizure duration. Recruitment of ANT was significantly higher for seizures that lasted longer (p 0.005), and the probability of recruiting ANT increased for longer seizures (Figures 5A and B). As biological processes like seizure dynamics can be non-linear, we tested the possibility of nonlinear interactions by normalizing the-a) recruitment latencies to the seizure duration; and b) the seizure duration by dividing with the maximal seizure duration within every patient. With these normalizations, two distinct groups of seizures emerged that were separated by a normalized recruitment latency of 0.3 (Figure 5C). The first group (normalized recruitment latency <0.3) consisted of seizures with impaired consciousness (FIAS, FBTCS) that correlated with longer seizure duration (r=-0.51, p=0.002). The second group (normalized latency >0.3) consisted of FAS and E seizures that correlated with shorter duration (−0.78, p=0.004). There was no significant difference between the slopes of the two correlations (Fisher r-to-z: z=1.34, p=0.18) (Figure 5C). There was a significant negative correlation between the thalamic recruitment latencies (L1) and overlap parameter, implying that early recruitment of ANT is associated with sustained neural activity coordinated between EZ and ANT (Figure S2-F).

**Figure 5.**
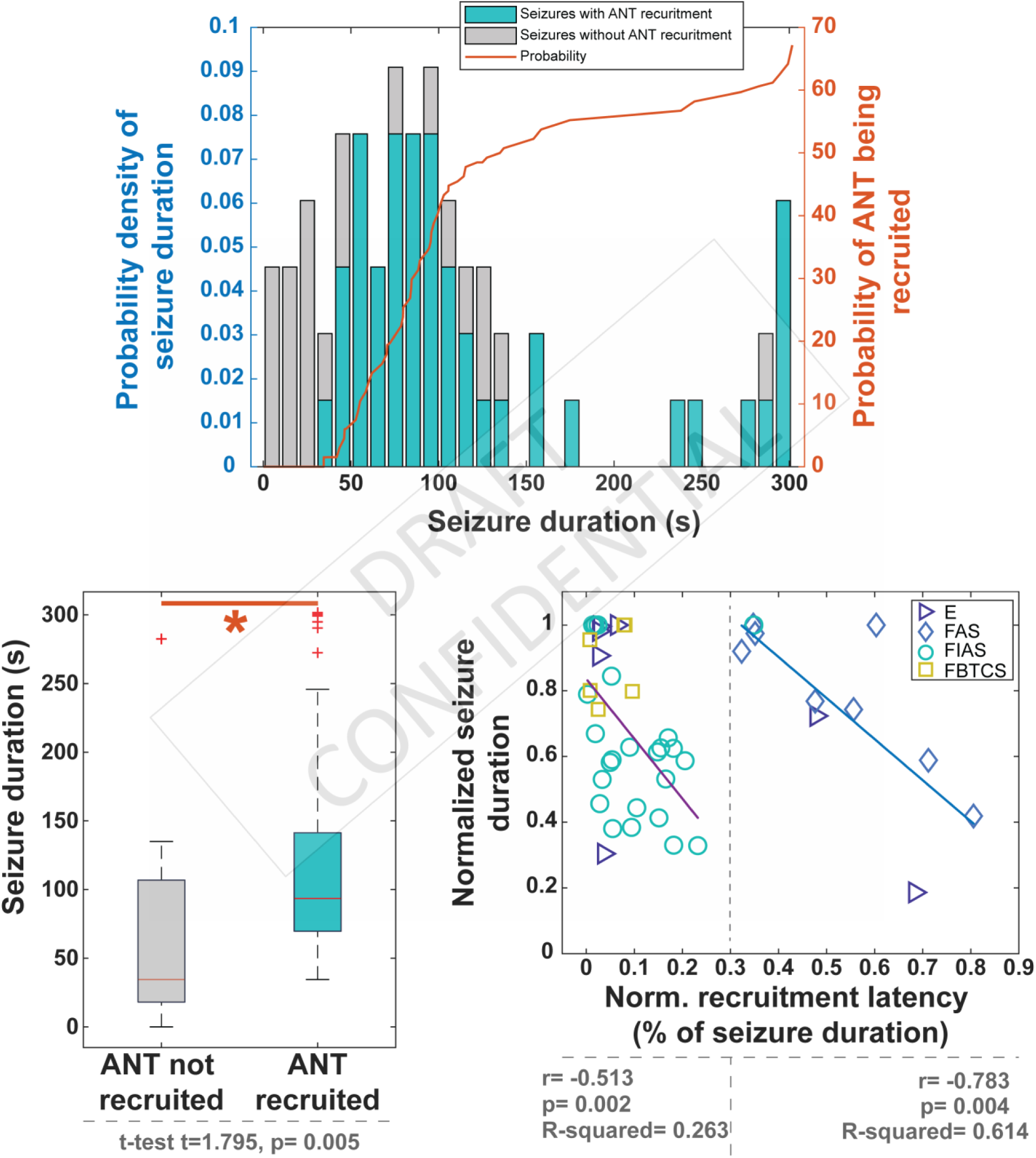
Anterior nucleus of thalamus (ANT) recruitment is dependent on seizure duration: A logistic regression analysis of the clinical predictors of ANT recruitment was performed and we found that the single most important predictor of recruitment was the duration of the seizure. (A) Shows the probability of ictal thalamic recruitment (orange axis on the right) increased with prolonged seizure duration. (B) T-test shows that longer seizures were associated with ANT recruitment. (C) It was noted that seizures tend to last longer when ANT was recruited faster. This was noted in both FIAs and FBTCS (r=-0.51, p=0.002) and FAS and ES (−0.78, p=0.004). There was no significant difference between the slopes of the two correlations (Fisher r-to-z: z=1.34, p=0.18).

### Spectral and MSE changes in the thalamogram during focal seizures

Following seizure onset, the first 12 seconds showed a progressive increase in power in theta, alpha and lower beta bands (4-16 Hz) for all seizure types except FBTCS and a decrease in the delta (1-2 Hz) and gamma bands (32-64 Hz) (Figure 6 A-E; Figure S1). The FBTCS seizures had increased 8-32 Hz band. Overall, the consistent spectral changes seen within the ANT during the early organization of seizure genesis were – a) increase in theta band power and b) a decrease in lower delta and gamma-band. The peak changes in theta band were seen around 26-31 seconds after seizure onset in the cortex. Interestingly these changes in spectral bands were also present for seizures that were not detected by LL, and the changes were independent of pre-seizure SOV (awake and sleep) (Figure 6 F-G). Following the seizure onset, there was a decrease in MSE measures in the ANT, thereby implying that the degree of randomness of thalamic LFP reduced during seizures. The changes were noted for all types of seizures (ES, FAS, FIAS and FBTCS)(Figure 6H).

**Figure 6:**
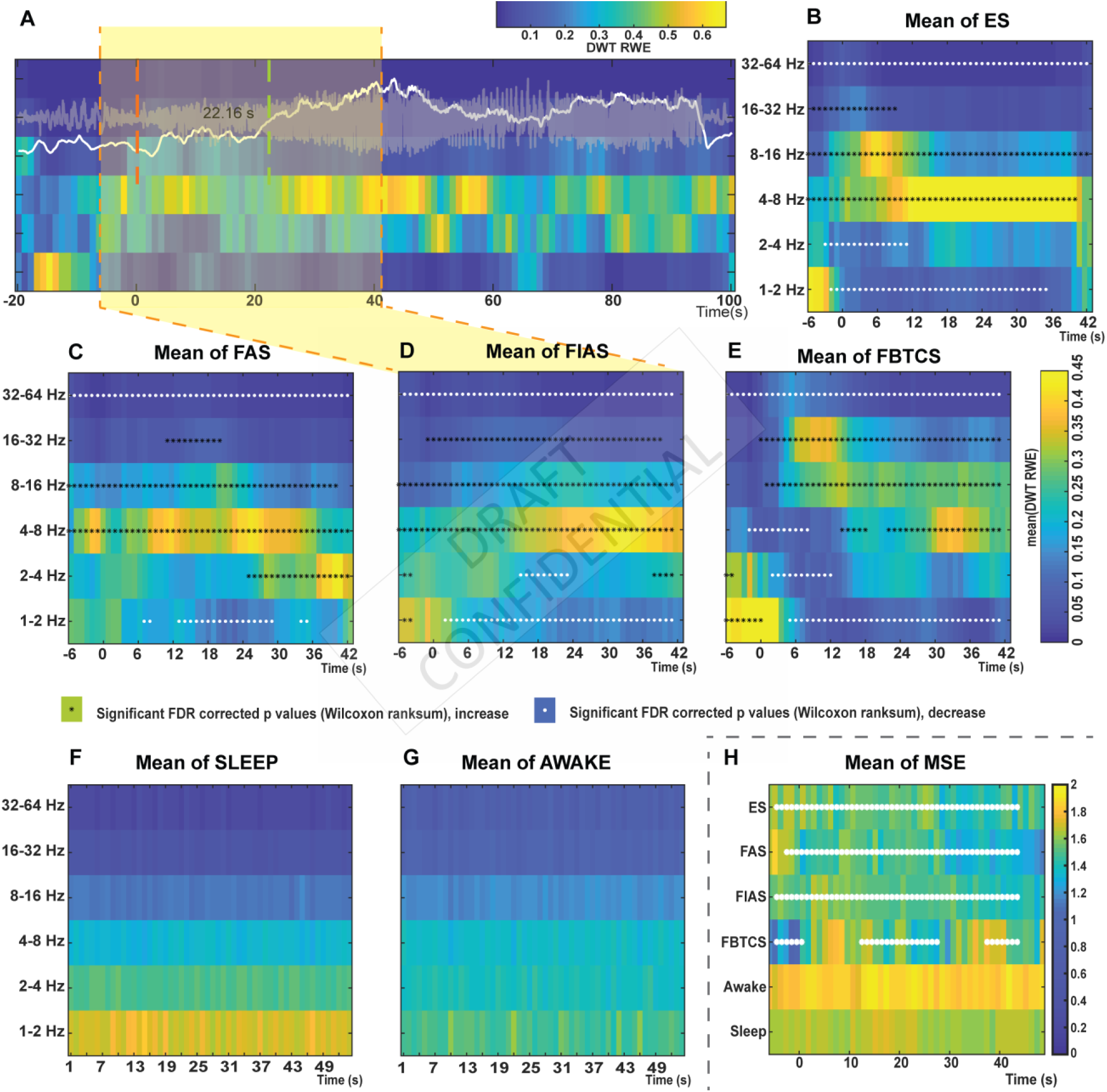
Temporal changes in discrete wavelet transform relative wavelet energy (DWT RWE) and multiscale entropy(MSE) across different ictal and interictal states recorded in the anterior nucleus of thalamus: The series of graphs try to explain spectral power changes (increases: black asterisks and decreases: white asterisks). Increase in spectral power was noted between 4-32Hz while decrease was noted majorly in 1-2Hz and 32-64Hz for all the 4 seizure types. This increase was compared with the baseline EEG activity and compared using Wilcoxon Rank Sum test. The comparison was made using a moving window method. The multiple comparisons were FDR corrected.

### Electrophysiological signatures of the thalamic ictal state are distinct from interictal states

Seizures can induce changes in behavioral states (like arousal or altered vigilance) that can be associated with changes in the oscillatory power at narrow frequency bands. To evaluate if the temporal and spectral features (MSE and DWT RWE) can distinguish the thalamic ictal state from other interictal states (awake and sleep), we used two machine learning algorithms-RF and RKS. Both RF and RKS were able to classify the thalamic ictal state from other interictal states with higher accuracy (92.8 and 95.1 %) and precision (93.1 and 96%). All the features (DWT RWE between 1-64 Hz and MSE) were relevant for the classification, but the increased power in the theta band (4-8 Hz) had the highest discrimination capability (Figure 7).

**Figure 7.**
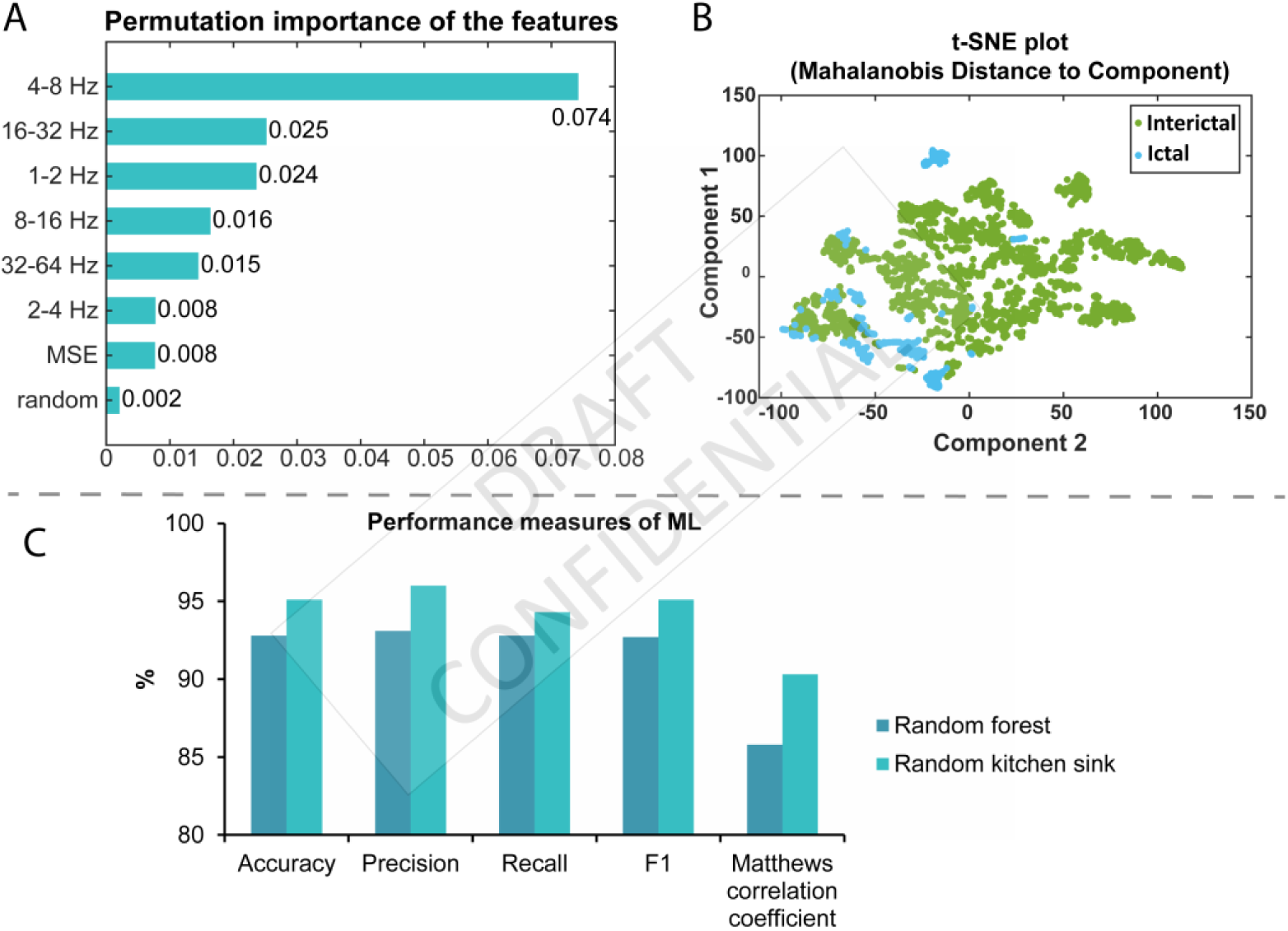
Spectral features (DWT-RWE and MSE) predict the ictal recruitment of anterior nucleus of thalamus(ANT) compared to background activity: Seven spectral features were used to predict the ictal recruitment of thalamus, i.e., the DWT-RWE measure of the 6 different frequencies and the MSE. (A) The bar plot shows the permutation importance of the 7 features. While the entire Random Forest model was largely predictive of distinguishing the ictal vs the interictal states, the DWT RWE of the theta and beta frequency bandwidths had the highest permutation importance. (B) A nonlinear dimensionality reduction using t-distributed stochastic neighbor embedding (t-SNE) shows that ictal DWT RWE and MSE cluster is distinct from interictal DWT RWE and MSE. (C) As an initial classification algorithm, random forest (RF) had an accuracy of 92.8% and a precision (positive predictive value) of 93.1%. As a validation we used a random kitchen sink (RKS) learning which also showed a higher accuracy (95.1%) and PPV of 96%. Overall these results point to the fact that spectral features of ANT recruitment can reliably predict ictal from interictal states. This implies that temporal lobe seizures are associated with distinct spectral changes in thalamus which possibly facilitate the myriad of clinical features in these patients.

### Replicating temporal patterns in ANT recruitment with stimulation-induced habitual seizures

Stimulation of ANT failed to trigger a seizure in all participants. Stimulation of EZ (amygdala, hippocampus) induced habitual seizures (3 FSA, 4 FIAS) in 7 participants. Interestingly the induced seizures recruited the ANT as was confirmed with line length and on visual inspection (Figure 8). The recruitment of ANT preceded the clinical onset for all the seizures (Figure 8).

**Figure 8.**
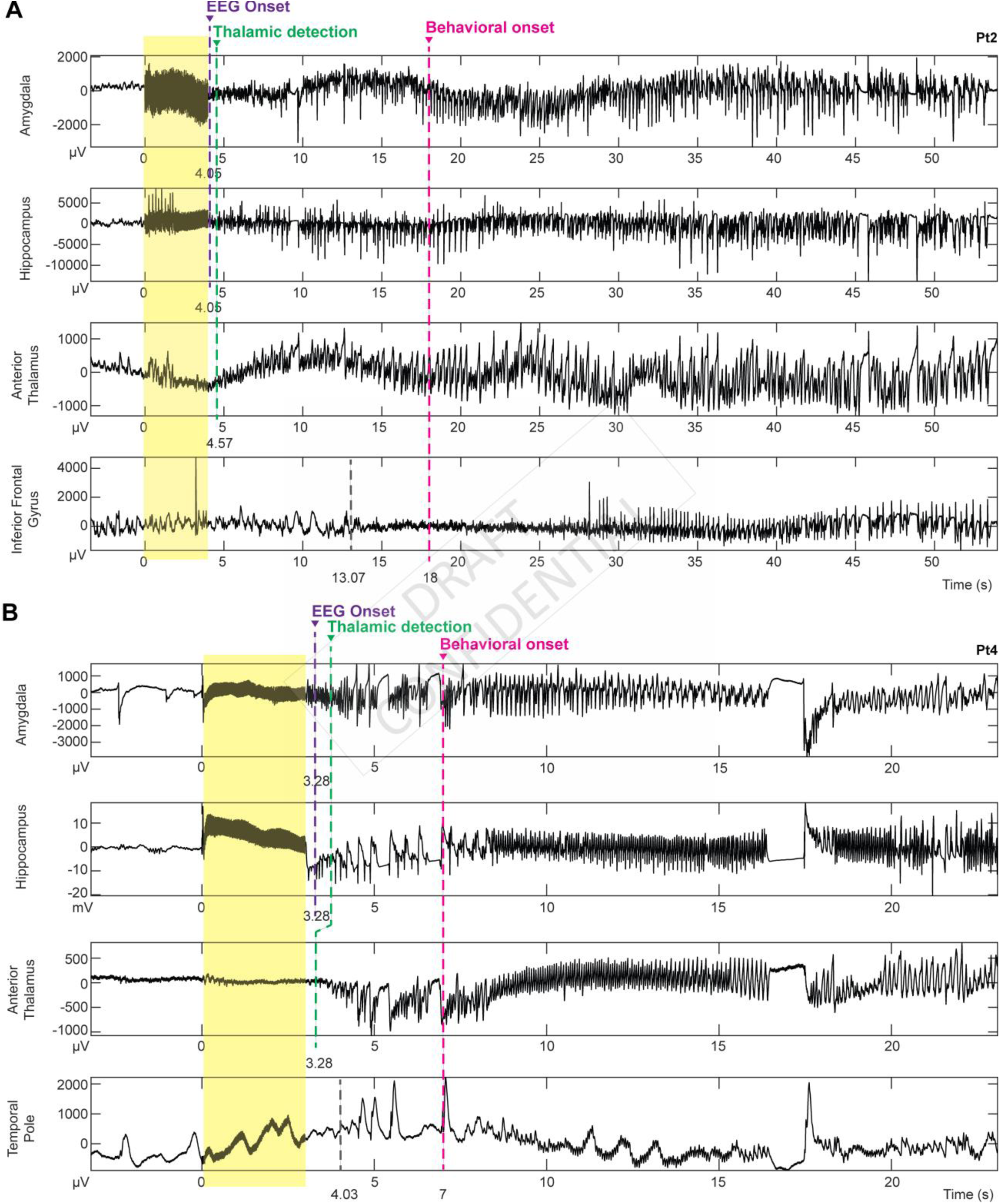
Electrical stimulation of epileptogenic zone induced seizures that recruited anterior nucleus of the thalamus (ANT): Two patients had stereotypical habitual electro-clinical seizures following stimulation of the EZ. We noted that the stimulation induced seizures recruited the ANT as evident by ictal EEG changes that were detected by line length (green dotted line). Behavioral manifestation of induced seizures (magenta dotted line) succeded ANT recruitment. The ictal EEG change in the EZ was also confirmed using Line Length (EZ detection – purple dotted line).

### Clinical predictors of thalamic recruitment in focal epilepsy

Seizure duration was the singular significant clinical predictor of ANT recruitment in this small cohort of suspected TLE (β=0.04, p=0.03). Higher seizure frequency, impairment of consciousness (FIAS, FBTCS), age of onset, duration of epilepsy, history of status epilepticus, MRI lesion, gender, and age were not found to be associated with thalamic recruitment in this cohort.

## Discussion

Rapid advancement in brain-computer interfaces and the long-term safety with invasive neuromodulation therapies have generated immense enthusiasm to develop adaptive or closed-loop interventions targeting the thalamocortical nodes in drug-resistant epilepsies (Osorio *et al*., 2015; Gummadavelli *et al*., 2018; Elder *et al*., 2019). However, the major impediment to the progress is the lack of peri-ictal electrophysiological recordings directly from the thalamocortical sites that might elucidate the underlying mechanisms and identify candidate biomarkers necessary for feedback stimulation. Capitalizing on the SEEG methodology that allowed recordings from multiple cortico-subcortical regions with over 1800 channels during 90-patient days (N=10), we have performed a comprehensive study to determine the patterns of ictal recruitment of ANT in TLE syndrome. Previous studies have utilized simultaneous recordings of EEG from the scalp and ANT DBS to study ictal recruitment of thalamus(Osorio *et al*., 2015; Krishna *et al*., 2016). Using SEEG that provides a superior temporospatial resolution than scalp EEG, we observe that –1) seizures localized to mesial temporal subregions (hippocampus, amygdala, anterior cingulate) have a higher predilection for ANT recruitment; 2) the recruitment is faster and remained sustained for seizures that impaired consciousness (FIAS and FBTCS), and 3) the temporal trends in the recruitment latencies is distinguished by seizure length. These recruitment patterns are consistent with preclinical studies that have demonstrated the role of ANT in propagation and sustaining of limbic seizures that progress to generalized tonic-clonic seizures (Mirski *et al*., 2003; Hamani *et al*., 2004; Takebayashi *et al*., 2007; Hamani *et al*., 2008; Feng *et al*., 2017). Lastly, 4) using data-driven analytics, we have identified temporo-spectral features that are provisional biosignatures of the thalamic ictal state. These results prompt a mechanistic role for the ANT in the early organization and sustaining of seizures, and the possibility to serve as a target for therapeutic closed-loop stimulation in TLE.

### Ictal recruitment of ANT depends on seizure onset sites

Underlying cortico-thalamic projections influence the ictal recruitment of thalamic subnuclei. The ANT receives afferents from the anterior cingulate cortex, hippocampus via fornix and mammillary bodies through the mammillothalamic tract(Child and Benarroch, 2013). Seizures can propagate to ANT directly from the epileptogenic regions that are localized within the limbic network or indirectly after propagating to regions whose afferents back project to the ANT. Our study confirms that the ANT is recruited by seizures that had at least one of the onset sites localized to hippocampus, amygdala, or anterior cingulate. Focal seizures that initiated and remained spatially confined to regions like the orbitofrontal and lateral prefrontal regions did not recruit ANT. Importantly, seizures were detected in the ANT in patients with multifocal (#2,7,10), and TLE plus epilepsies (#1, 5) when the ictal onset network included at least one of the limbic structures (hippocampus or anterior cingulate or amygdala).

### Recruitment of ANT varies with seizure duration and types

The ANT mediates cortical-subcortical interactions between the limbic system and the brainstem. These pathways are associated with a bihemispheric propagation of convulsive seizures(Norden and Blumenfeld, 2002). Perturbation of ANT by lesioning or high-frequency stimulation disrupted the seizure progression in preclinical models, thereby establishing a causal role of ANT in propagation(Hamani *et al*., 2004; Takebayashi *et al*., 2007). The fast recruitment of ANT in FIAS and FBTCS (Figures 4 and 5) support a growing body of evidence that suggests that the network dynamics at seizure onset and early propagation can vary with seizure types(Karoly *et al*., 2018; Pizarro *et al*., In-Press). Insights from the dynamics of seizure initiation and termination suggest that the state transition from a seizure to the interictal state is not a random fluctuation in cortical activity(Kramer *et al*., 2012; Jirsa *et al*., 2014; Cook *et al*., 2016; Bauer *et al*., 2017). The seizure follows a set pathway that must be completed before entering the interictal state. Cook et al. showed that seizure duration is bimodally distributed with different types have a distinct onset and offset mechanism(Cook *et al*., 2016). The observation agrees with our results that show seizure length varied with ANT recruitment latencies. The presence of sustained thalamocortical interactions in FIAS and FBTCS, as evidenced by the overlap duration, provides an opportunity to intervene and disrupt the propagation of those seizures that are associated with higher morbidity and mortality(Thurman *et al*., 2017).

### States of vigilance influence thalamic detection of seizures

Epilepsy and sleep have a complex bidirectional relationship. Sleep can affect seizure occurrence and seizure threshold. The thalamus is intricately related to the modulation of the sleep-wake cycle and, hence, understanding thalamic recruitment during different SOV is of great importance for developing neuromodulation strategies(Llinas and Steriade, 2006; Sedigh-Sarvestani *et al*., 2014; Ewell *et al*., 2015). As anticipated, our study demonstrated faster propagation of seizures to the thalamus during sleep. This finding may be important in the context of previous studies that documented generalization of temporal lobe seizures to be more frequent during sleep. The opportunity to rapidly detect seizures in the thalamus allows the implementation of a feedback stimulation early at the onset that might abort seizure progression.

### Thalamic fingerprints of mesial temporal lobe seizures

Seizure related electrographic and spectral changes in the cortex are well established, but in the thalamus, these changes are not well defined. Therefore, we have adopted multiple methods (EI, LL, wavelet and entropy-based methods) that are based on different principles to detect a seizure. The frequency parameters to calculate EI was between 12-127 Hz/4-12 Hz. Note, for the EI to be higher than 0.2, either the frequency contents of the field potentials need to be higher than 12 Hz, or the bandwidth 4-12 Hz need to decrease at seizure onset. The spectral analysis demonstrated that thalamic ictal state was associated with a fast decrease in gamma and delta and a slow increase in the theta. Therefore, not surprisingly, only 21% of seizures (in five subjects) had higher EI. The LL detection, which is based on variation in signal amplitude and frequency, performed slightly better than EI but still failed to detect or had long latency in detecting seizures that were associated with early changes in theta bandwidth. Wavelet-based methods can detect the thalamic ictal state transitions that were predominantly in lower frequencies, and the methods may be an attractive solution for its ability to compute fast that is necessary for any feedback interventions.

### The emergence of theta rhythms in the ANT during focal seizures

Although multiple spectral changes characterize the thalamic ictal state, the emergence of theta rhythm (4-8 Hz) was significantly higher and sustained as seizures evolved. Vertes et al. and Tsanov et al. demonstrated the presence of theta-bursting neurons in the anterior thalamus that was entrained to the Papez circuit (including the medial septum, hippocampus, and ANT)(Vertes *et al*., 2001; Tsanov *et al*., 2011). We speculate that the theta rhythm integrates functionally the ANT in the limbic-diencephalic circuit with the progression of mesial temporal lobe seizures. The endogenous state that is characterized by theta rhythm is associated with the increased cholinergic activity, seizure severity, and has been found to either promote or prevent a seizure(Colom *et al*., 2006; Sedigh-Sarvestani *et al*., 2014; Ewell *et al*., 2015; Yi *et al*., 2015). Electrical stimulation in theta frequency ameliorated epileptic discharges in preclinical models of limbic epilepsy(Sedigh-Sarvestani *et al*., 2014). Further work is necessary to establish the role of theta rhythm in thalamo-hippocampal seizures, especially in preventing generalization of limbic seizures.

### Study limitation

The goal of the study was to understand the ANT recruitment patterns in mesial TLE, and hence, we selected patients with suspected TLE who are undergoing SEEG investigation. However, the post-SEEG localization of EZ in some patients extended beyond the amygdala-hippocampus to extratemporal regions, and this heterogeneity in cohort contributed to variability in recruitment patterns. Thalamus is not implanted regularly during SEEG investigation, and hence our cohort was limited to 10 subjects. Anti-seizure drugs (ASD) are tapered off routinely in the epilepsy monitoring, and this may confound the recruitment patterns by facilitating seizure propagation. However, the stimulation-induced seizures, that were performed after reloading the ASD, were able to replicate the temporal trends in recruitment seen with the spontaneous seizures.

## Conclusions

Temporal lobe seizures can recruit the anterior nucleus of the thalamus, but details about recruitment patterns were unknown. We confirm that seizures localized to mesial temporal subregions (hippocampus, amygdala, anterior cingulate) can recruit ANT early before the clinical manifestation. Recruitment latencies are influenced by seizure types, pre-seizure SOV, and seizure length. The thalamic ictal state is characterized by specific temporo-spectral changes that are candidate biosignatures for detecting focal seizures. Overall, our results prompt a mechanistic role for the ANT in the early organization of mesial temporal seizures and suggest that field potentials recorded from ANT can be targeted for therapeutic closed-loop interventions.

## Abbreviations

ASD: Anti-seizure drugs
DWT: Discrete Wavelet Transfer
EI: Epileptogenecity Index
ES: Electrographic Seizures
EZ: Epileptogenic Zone
FAS: Focal Onset Aware Seizures
FBTCS: Focal to bilateral tonic-clonic seizures
FIAS: Focal onset seizures with impaired awareness
HAAC: Hippocampus, Amygdala, Anterior Cingulate complex
LL: Line Length
MSE: Multiscale Entropy
RF: Random Forest
RKS: Random Kitchen Sink
RWE: Relative Wavelet Energy
SEEG: stereo-electroencephalography
TLE: Temporal lobe epilepsy
UEO: Unequivocal electrographic onset

## Acknowledgments

We would like to acknowledge the contribution of UAB Surgical Epilepsy team and patients who participated in the research with the hope of contributing to a science that is focused on developing a therapy to improve outcome.

